# Loss of PTPRK in hepatocytes reduces steatosis and carcinogen-induced tumour development in obesity

**DOI:** 10.64898/2026.01.17.697695

**Authors:** Ao Li, Wadsen St-Pierre-Wijckmans, Gabriel G. Hovhannisyan, Tiffany Lai, Carlos E. Buss, Erick Arroba, Rabéa Dahili, Leila Hosseinzadeh, Sumeet Pal Singh, Lu Yu, David Oxley, Hayley J. Sharpe, Eduardo H. Gilglioni, Esteban N. Gurzov

**Affiliations:** Signal Transduction and Metabolism Laboratory, Faculty of Medicine, Université libre de Bruxelles, Brussels B-1070, Belgium; Regenerative Biology Lab, Institut de Recherche Interdisciplinaire en Biologie Humaine et Moléculaire (IRIBHM), Université libre de Bruxelles, Brussels B-1070, Belgium; Signalling Programme, Babraham Institute, Babraham Research Campus, Cambridge CB22 3AT, United Kingdom; Department of Life Sciences, School of Natural Sciences, Shiv Nadar Institution of Eminence, Delhi 201314, India; Proteomics, The Babraham Institute, Cambridge, CB22 3AT, United Kingdom

## Abstract

Protein tyrosine phosphatases are crucial regulators of metabolism with specific roles in different tissues. To investigate hepatocyte-specific function of protein tyrosine phosphatase receptor type K (PTPRK), we generated mice carrying floxed *Ptprk* alleles and crossed them with Alb-Cre mice (*Ptprk^ΔHep^* mice). Under chow feeding, *Ptprk^ΔHep^*mice were largely comparable to littermate controls. In contrast, *Ptprk^ΔHep^*mice fed a high-fat, high-fructose, high-cholesterol diet exhibited reduced steatosis, lower hepatic PPARγ, and blunted hepatocyte hypertrophy, accompanied by improved systemic insulin sensitivity, as assessed by hyperinsulinemic-euglycemic clamps. We identified PTPRK-interacting proteins enriched for metabolic functions associated with glycolysis and lipid biosynthesis using pull downs from primary hepatocyte lysates. In line with these findings, *Ptprk^ΔHep^* mice developed fewer tumours than controls in an obesity and carcinogen-induced hepatocellular carcinoma (HCC) model. Our data show that under nutrient excess PTPRK is functionally engaged in hepatocytes to support PPARγ-linked steatotic growth, insulin resistance, and tumour initiation, highlighting PTPRK as a potential therapeutic target in MASLD-associated HCC.

## Introduction

Excessive caloric intake associated with modern lifestyles represents a major challenge for human health. The global rise in obesity has driven parallel epidemics of metabolic dysfunction-associated steatotic liver disease (MASLD) and hepatocellular carcinoma (HCC),^1,2^ with adult MASLD prevalence predicted to increase within the next 15 years.^3^ Hepatocytes adapt to nutrient excess by reprogramming their metabolism to optimise energy storage pathways, resulting in increases in cell size that accommodate accumulated glycogen and triglycerides.^4^ Esterification of fatty acids into triglycerides and their storage within lipid droplets constitutes a protective mechanism that limits free fatty acid toxicity and buffers acute lipid overload in the circulation. Similarly, spare carbohydrates that exceed glycogen storage capacity are redirected toward lipogenesis. While these responses are initially adaptive, chronic overnutrition drives sustained metabolic reprogramming, insulin resistance, and a markedly increased risk of hepatocarcinogenesis.^4–6^ Although lifestyle interventions remain first-line therapies, a deeper molecular understanding of how hepatocytes sense, integrate, and adapt to nutrient overload is required to identify additional therapeutic targets.^4,7,8^

Protein tyrosine phosphorylation is a central mechanism through which hepatocytes dynamically coordinate metabolic flux in response to nutritional and hormonal cues.^9,10^ Receptor-linked protein tyrosine phosphatases can integrate extracellular signals with intracellular metabolic control, yet their specific roles in hepatocyte metabolism remain incompletely understood. Protein Tyrosine Phosphatase Receptor Kappa (PTPRK) has recently emerged as an important regulator of hepatic glycolysis and lipid metabolism.^11^ PTPRK is a receptor-type phosphatase that localises to intercellular junctions and engages in homophilic interactions with adjacent cells via its extracellular domain.^12^ PTPRK-deficient mice (*Ptprk^−/−^*) do not exhibit overt metabolic abnormalities under chow-fed conditions but display delayed progression of diet-induced obesity-associated complications, including hepatic steatosis, insulin resistance, and HCC.^11^ PTPRK promotes glycolysis-mediated PPARγ expression in the liver, promoting *de novo* lipogenesis, and linking glucose handling to lipid synthesis under nutrient-rich conditions.^11,13^

Context-dependent functions were reported for PTPRK across different tissues. In the intestinal epithelium, PTPRK maintains proliferative homeostasis by negatively regulating EGFR-ERK signalling, and its loss leads to enterocyte hyperproliferation, crypt hyperplasia, and impaired nutrient absorption in celiac disease models.^14,15^ In vascular endothelial cells, PTPRK regulates angiogenesis and contributes to pathological vascular remodelling during metabolic inflammation and tumourigenesis.^16^ PTPRK is also expressed in the central nervous system, where it functions as a regulator of cell-cell adhesion and neuronal circuit assembly, contributing to axon guidance, synaptic targeting, and neuronal connectivity through selective modulation of signalling pathways during neural development.^17^ Collectively, these observations indicate that PTPRK acts as a versatile, tissue-specific regulator whose functional output is dictated by cellular context and signalling environment.

Our previous work using whole-body *Ptprk* inactivation established PTPRK as a regulator of obesity-driven hepatic glycolysis, PPARγ signalling, steatosis, and tumour susceptibility.^11^ It remained unclear whether PTPRK deficiency specifically in hepatocytes is sufficient to drive metabolic adaptations to lipid overload and protect against liver disease progression. Hence, we developed a novel conditional *Ptprk^Lox/Lox^* mouse model and generated hepatocyte-specific PTPRK knockout mice (*Ptprk^ΔHep^*). Our findings identified hepatocyte-intrinsic PTPRK as a key regulator of nutrient-driven lipid accumulation and metabolic stress in obesity, with potential implications for liver cancer prevention in the context of obesity-associated MASLD.

## Results

### *Ptprk^ΔHep^* attenuates liver fat accumulation, hepatic PPARγ expression, and systemic insulin resistance in diet-induced obese mice

We generated a novel *Ptprk^Lox/Lox^* mouse model in which exon 3 of *Ptprk* is flanked by loxP sites using CRISPR/Cas9 genome editing. Hepatocyte-specific deletion was achieved by crossing *Ptprk^Lox/Lox^* mice with Alb-Cre mice^18^ on a C57BL/6N background, resulting in the *Ptprk^ΔHep^*mouse line (**Fig. 1a**).

**Fig. 1.**
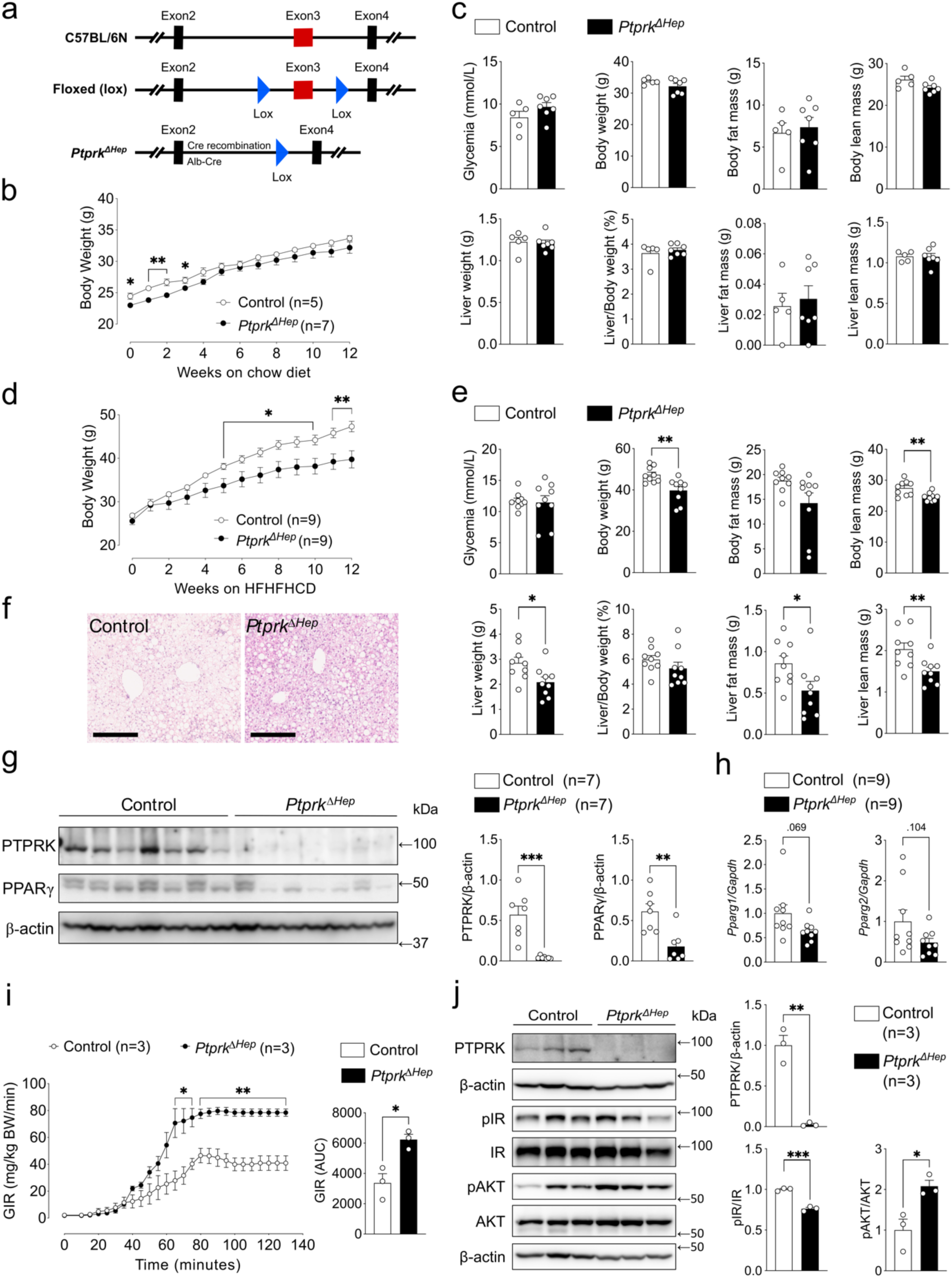
Hepatocyte-specific *Ptprk* deletion (*Ptprk^ΔHep^*) protects against diet-induced obesity, hepatic fat accumulation, and systemic insulin resistance. (a) Schematic of hepatocyte-specific *Ptprk* knockout (*Ptprk^ΔHep^*) or littermate control (*Ptprk^Lox/Lox^*) and mice. (b,c) Body weight progression (b) and metabolic parameters (c) in 8 weeks old *Ptprk^ΔHep^*(n=7) and control (n=5) mice fed a chow for additional 12 weeks. (d,e) Body weight progression (d) and metabolic parameters (e) in *Ptprk^ΔHep^* (n=9) and control (n=9) mice starting at 8 weeks of age a high-fat, high-fructose, high-cholesterol diet (HFHFHCD) for 12 weeks. (f) Representative haematoxylin and eosin staining of liver sections from *Ptprk^ΔHep^* and control mice at week 12 of HFHFHCD feeding. Scale bar = 250μm. (g,h) Immunoblot (n=7) (g) and RT-PCR (n=9) (h) analyses of liver tissues from *Ptprk^ΔHep^* and control mice. (i,j) Hyperinsulinemic-euglycemic clamp (i) and corresponding immunoblot (j) analysis showing glucose infusion rate (GIR) and hepatic protein levels of PTPRK, pIR, and pAKT in *Ptprk^ΔHep^* (n=3) and control (n=3) mice fed with HFHFHCD for 6 weeks (14-week-old). Data are presented as mean ± SEM (b-e, g-j). Statistical significance was performed using two-tailed unpaired Student’s *t*-tests (b-e, g-j). \**p* < 0.05, \*\**p* < 0.01, and \*\*\**p* < 0.001.

To assess the metabolic effects of hepatocyte-specific PTPRK deletion, we monitored body weight, body composition, and liver parameters in *Ptprk^ΔHep^* and control littermates maintained on a chow diet (**Fig. 1b**). Both groups showed progressive weight gain beginning at 8 weeks of age, with *Ptprk^ΔHep^* males maintaining body weights with slight differences compared to littermate controls. After 12 weeks of chow diet feeding, *Ptprk^ΔHep^* mice exhibited no significant differences in glycemia (9.67 ± 0.5 vs. 8.42 ± 0.73 mmol/L), body fat mass (7.37 ± 1.16 vs. 6.68 ± 1.2 g), or liver fat mass (∼0.03 ± 0.01 g in both groups) when compared with controls (**Fig. 1c**). Other parameters, including total body weight, lean mass, liver weight, liver-to-body weight ratio, and liver lean mass, were also unchanged between groups (**Fig. 1c**). Immunoblot analysis of liver lysates showed no significant difference in PPARγ protein expression between *Ptprk^ΔHep^* and control mice (**Supplementary Fig. 1a**). Collectively, these findings indicate that hepatocyte-specific PTPRK deletion does not significantly affect metabolic parameters or hepatic PPARγ expression under chow diet-fed conditions.

We next assessed *Ptprk^ΔHep^* and control mice fed a high-fat, high fructose, high cholesterol diet (HFHFHCD) starting at 8 weeks of age (**Fig. 1d**). *Ptprk^ΔHep^* mice gained weight more gradually than controls, leading to significantly lower body weights during weeks 5–12. After 6 weeks of HFHFHCD feeding, *Ptprk^ΔHep^* mice displayed reduced body weight (35.19 ± 1.76 vs. 39.76 ± 0.84 g) and lean mass (23.12 ± 0.5 vs. 25.25 ± 0.69 g) compared to controls (**Supplementary Fig. 1b**). At week 12, while glycemia remained unchanged between groups (11.46 ± 1.1 vs. 11.72 ± 0.44 mmol/L), *Ptprk^ΔHep^* mice continued to show lower body weight, body lean mass, liver weight, liver fat mass, and liver lean mass when compared with controls (**Fig. 1e**).

To evaluate whole body metabolic function, we performed indirect calorimetry analysis at week 12 using metabolic cages. No significant differences were observed in water and food intake, energy expenditure, respiratory exchange ratio, oxygen consumption, or ambulatory activity between *Ptprk^ΔHep^*and control mice (**Supplementary Fig. 1c**). Histological assessment using haematoxylin-eosin (H&E) staining revealed visibly reduced hepatic steatosis in *Ptprk^ΔHep^* mice (**Fig. 1f**). Consistently, immunoblot analysis showed a marked reduction in hepatic PPARγ protein levels (**Fig. 1g**), and a downregulation trend of *Pparg* mRNA (**Fig. 1h**) in *Ptprk^ΔHep^* mice compared with littermate controls.

To directly assess insulin sensitivity, we performed hyperinsulinemic–euglycemic clamps in 14-week-old male *Ptprk^ΔHep^* mice after 6 weeks of HFHFHCD feeding. Basal and clamp glycemia were maintained at comparable levels between *Ptprk^ΔHep^* mice and controls (control area under the curve (AUC): 1028.1 ± 133; *Ptprk^ΔHep^* AUC: 943 ± 25) (**Fig. 1i, Supplementary Fig. 1d**), confirming equivalent glycaemic conditions. In contrast, *Ptprk^ΔHep^* mice required a significantly higher glucose infusion rate (GIR) to maintain euglycemia (GIR AUC: 6239.67 ± 343.54) compared with control littermates (GIR AUC: 3372 ± 609.13), indicating enhanced insulin sensitivity. Immunoblot analysis revealed reduced insulin receptor phosphorylation but markedly increased AKT phosphorylation at Ser473 in livers of *Ptprk^ΔHep^* mice relative to controls (**Fig. 1j**). These data indicate that hepatocyte PTPRK contributes to the development of insulin resistance under obesogenic conditions.^11^ Collectively, hepatic PTPRK deletion reduces hepatic lipid accumulation and insulin resistance under obesogenic conditions, likely via suppression of PPARγ.

### PTPRK engages metabolic control networks in primary mouse hepatocytes

Intracellular lipid droplets can mechanically alter hepatocyte size and impair insulin sensitivity.^19,20^ To assess the diet-dependent effects on hepatic morphology, we measured hepatocyte size in liver sections from *Ptprk^ΔHep^* and control mice after six weeks on either a chow diet or a HFHFHCD, using phalloidin staining (**Fig. 2a**). Under chow diet, hepatocyte morphology and quantified cell area were comparable between control (318.01 ± 7.61 μm²) and *Ptprk^ΔHep^* (328.12 ± 8.22 μm²) mice (**Fig. 2a**). HFHFHCD increased the size of control hepatocytes (406.32 ± 19.11 μm²), which was prevented in *Ptprk^ΔHep^* hepatocytes (327.82 ± 14.15 μm²) (**Fig. 2a**).

**Fig. 2.**
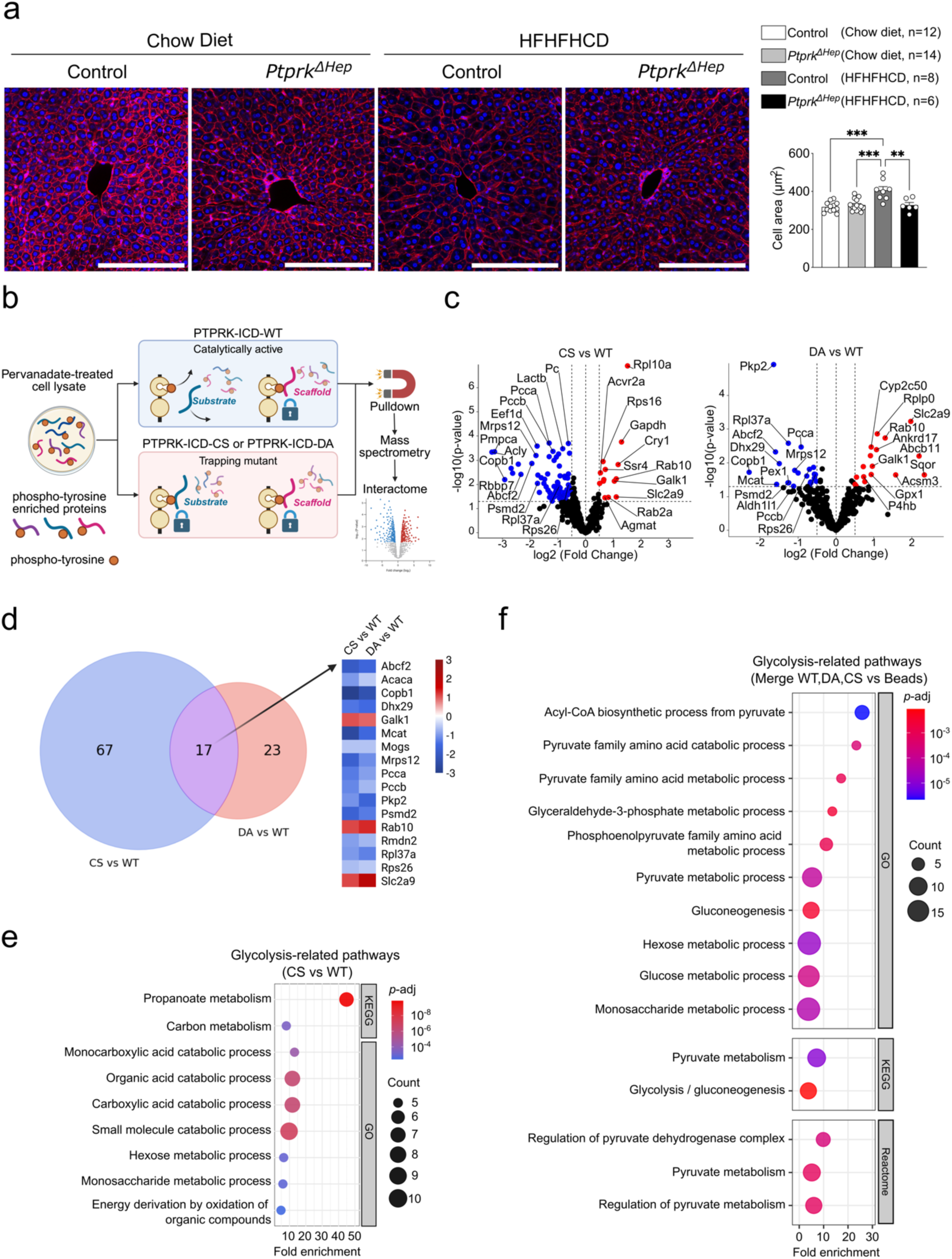
*Ptprk^ΔHep^*reduces hepatocyte size, and PTPRK ICD mutation alters glycolysis-related pathways. (a) Liver phalloidin staining in *Ptprk^ΔHep^* and control mice fed chow diet or HFHFHCD for 6 weeks (age 14 weeks) (n=6-14). Scale bar = 200μm. (b) Schematic overview of the interactome experiment with different PTPRK protein variants. (c) Volcano plots of differently enriched p-tyrosine pull-down proteins in CS vs WT (left) and DA vs WT (right). (d) Venn diagram of significantly enriched p-tyrosine pulldown proteins in CS vs WT and DA vs WT. (e) ORA pathway enrichment of interactome showing glycolysis-related pathways from KEGG and GO in CS vs WT. (f) ORA pathway enrichment of interactome showing glycolysis-related pathways from GO, KEGG, and Reactome using all enriched proteins (WT vs Beads, DA vs Beads, CS vs Beads). Data are presented as mean ± SEM (a). Statistical significance was performed using one-way ANOVA (a). \*\**p* < 0.01, and \*\*\**p* < 0.001. In (e,f), FDR < 0.05.

To gain molecular insights into how PTPRK regulates hepatocyte function, we mapped the phosphotyrosine-associated interactome of recombinant PTPRK intracellular domain. Pull downs were carried out following incubation of recombinant PTPRK intracellular domains (ICD) with lysates from pervanadate-treated primary wild-type mouse hepatocytes and analysed by mass spectrometry (**Fig. 2b**). We compared proteins associated with the catalytically active wild-type ICD (WT) and two catalytically inactive mutants that act as substrate traps by disrupting distinct steps of the phosphatase reaction cycle: a Cys-to-Ser mutant (CS), which prevents nucleophilic attack by the catalytic cysteine, and an Asp-to-Ala mutant (DA), which blocks general acid-mediated bond cleavage and stabilises a closed catalytic pocket conformation (**Fig. 2b, Supplementary Fig. 3a-d**).

The intersection of WT, CS, and DA datasets yielded 25 proteins consistently present across all three conditions, but absent from the beads-only control (**Supplementary Fig. 3e**). We hypothesise that these represent scaffolded assemblies rather than direct substrates. We next focused on proteins specifically enriched by the substrate trapping mutants. There were unique and overlapping protein interactors for the two mutants, reflecting the distinct trapping properties of the two mutants. We identified 14 proteins significantly enriched by the CS mutant compared to the WT, and 17 proteins enriched by the DA mutant (**Fig. 2c**). Amongst these proteins, 3 were commonly enriched in both mutants (**Fig. 2d**). The proteins that showed increased abundance in both CS- and DA-trapping mutants relative to WT, and therefore candidate substrates, are the vesicular trafficking regulator Rab10, the carbohydrate metabolism enzyme Galk1, and the solute carrier Slc2a9 (**Fig. 2d**).

To explore functional pathways associated with CS-enriched proteins compared to WT, we performed over-representation analysis (ORA) using Gene Ontology (GO), Kyoto Encyclopedia of Genes and Genomes (KEGG), and Reactome annotations (**Supplementary Fig. 2f-h**) and also for all proteins enriched in WT, CS, and DA (**Supplementary Fig. 3a-c**). These analyses revealed enrichment across mitochondria-related functions, mitochondrial translation, fatty acid metabolism, acyl-CoA-linked processes, ribosome-associated pathways, and amino acid degradation, indicating broad engagement of PTPRK-associated networks with metabolic and translational machinery. Targeted, hypothesis-driven selection of pathways related to glycolysis is shown in **Fig. 2e** for CS vs WT groups and in **Fig. 2f** for the merged WT, CS, and DA datasets. Within this selected subset, CS-enriched proteins were associated with propanoate metabolism, carbon metabolism, carboxylic acid catabolism, hexose metabolic process, monosaccharide metabolic process, and energy derivation by oxidation of organic compounds (**Fig. 2e**). Integration of significantly enriched proteins across WT, CS, and DA conditions followed by selective examination of glycolysis-related annotations revealed pyruvate-centred processes (**Fig. 2f**). These included glycolysis and gluconeogenesis, pyruvate metabolism, regulation of pyruvate metabolism, and regulation of the pyruvate dehydrogenase complex, alongside upstream glucose, hexose, and monosaccharide metabolic processes. Although these pathways represent a curated subset of the broader enrichment landscape, their convergence across multiple comparisons positions PTPRK-associated protein networks at metabolic branch points that govern carbon flux toward mitochondrial oxidation versus anabolic lipid synthesis.

### *Ptprk^ΔHep^* reduces tumour multiplicity in obesity-associated HCC

PTPRK expression correlates with increased glycolytic and lipogenic gene expression during human HCC development.^11^ In *Ptprk^-/-^* mice, we previously observed reduced body weight, hepatic fat mass, and significantly attenuated DEN-induced tumourigenesis.^11^ To delineate the hepatocyte-specific contribution of PTPRK to HCC development, we induced liver cancer in *Ptprk^ΔHep^* and control mice using dimethylnitrosamine (DEN) under HFHFHCD for 25 weeks (**Fig. 3a**). *Ptprk^ΔHep^*mice exhibited similar body weight gain as controls (**Fig. 3b**. Moreover, among the DEN-injected mice used for metabolic and morphological assessments, no significant differences were observed between *Ptprk^ΔHep^* and control groups in glycemia (9.06 ± 0.69 vs. 9.88 ± 0.52 mmol/L), body weight, spleen weight, liver weight, liver-to-body weight ratio, fat liver mass, or average tumour size (2.82 ± 0.64 vs. 2.25 ± 0.12 mm) (**Fig. 3d**).

**Fig. 3.**
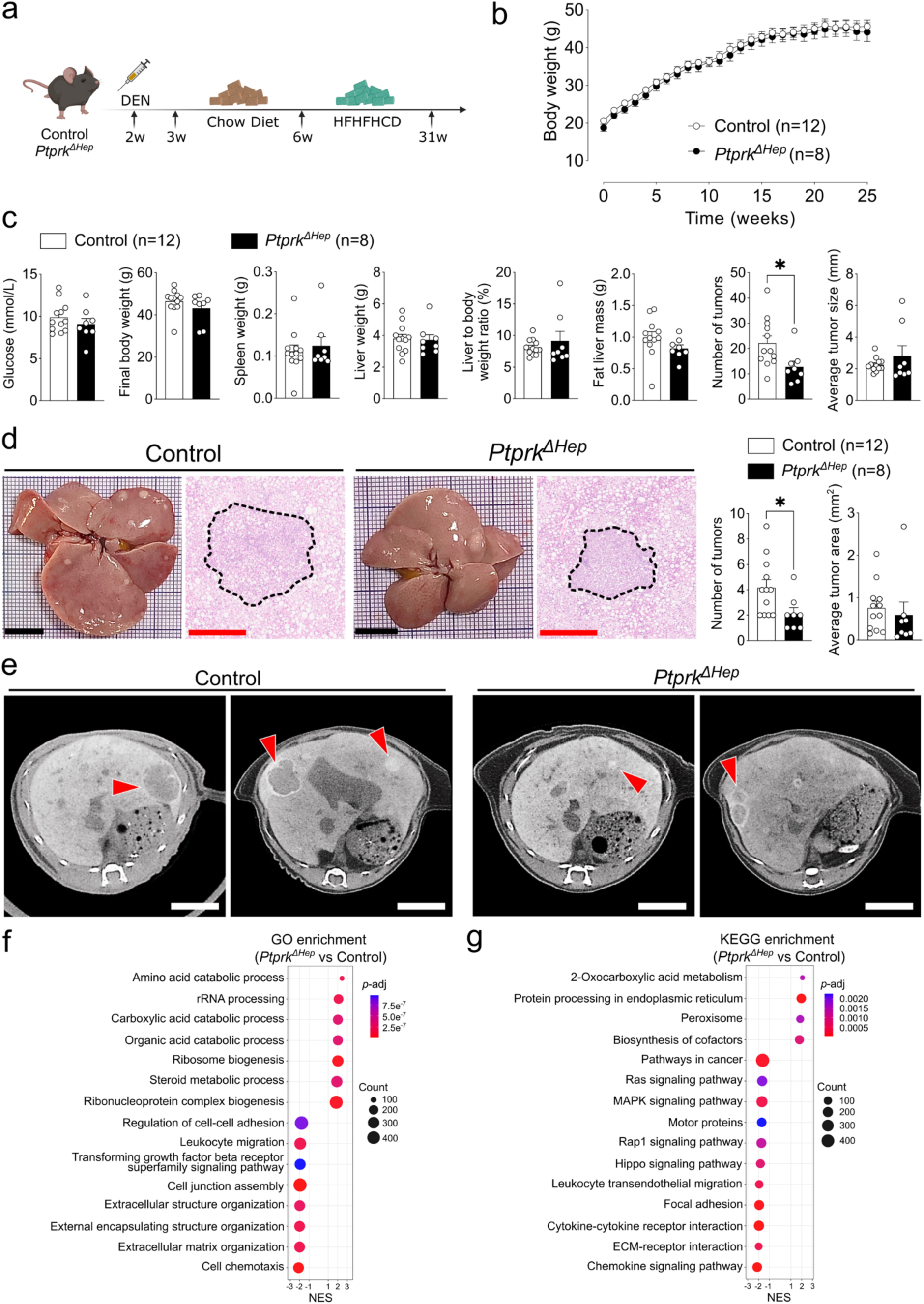
*Ptprk^ΔHep^* reduces tumour multiplicity in obesity-associated HCC. (a) Schematic overview of the DEN + HFHFHCD-induced HCC model in *Ptprk^ΔHep^* and control mice. (b) Body weight progression of DEN-injected *Ptprk^ΔHep^* (n=8) and control (n=12) mice fed HFHFHCD for 25 weeks. (c) Metabolic and tumour-related parameters of *Ptprk^ΔHep^* (n=8) and control (n=12) mice. (d) Histological analysis of liver sections stained with haematoxylin and eosin in *Ptprk^ΔHep^* (n=8) and control (n=12) mice. Black scale bar = 1cm. Red scale bar = 500μm. (e) Representative CT scan images of *Ptprk^ΔHep^*and control mice. Scale bar = 7mm. (f,g) GO (f) and KEGG (g) GSEA pathway enrichment analyses of transcriptomics in tumour samples comparing *Ptprk^ΔHep^* vs control. Data are presented as mean ± SEM (b,d,e). Statistical analyses were performed using two-tailed unpaired Student’s *t*-test (b,d,e). \**p* < 0.05. In (f,g), FDR < 0.05.

*Ptprk^ΔHep^* mice developed a reduced number of tumours (12.75 ± 2.31) compared to controls (22.17 ± 2.8) (**Fig. 3c**). Histological examination of liver sections showed no significant difference in average tumour area (0.59 ± 0.4 vs. 0.76 ± 0.11 mm^2^), but a reduced tumour numbers in *Ptprk^ΔHep^* mice (2.13 ± 0.31 vs. 4.17 ± 0.72) (**Fig. 3d**). Computed tomography (CT) imaging showed classic features of HCC in both groups, circumscribed hypodense masses and peripheral arc-like hyperattenuating fibrosis. However, *Ptprk^ΔHep^* livers showed visibly reduced hypodensity area and peripheral fibrosis, suggesting decreased tumour burden (**Fig. 3e**). Three-dimensional reconstruction further revealed that *Ptprk^ΔHep^* mice exhibited fewer tumours than control mice (**Supplementary Fig. 4a**).

We further performed bulk RNA sequencing on liver tumour samples from control and *Ptprk^ΔHep^* mice. The cell proliferation related gene *Wnt7a* and the angiogenesis related gene *Olfml2b* showed reduced expression in *Ptprk^ΔHep^* liver tumours, whereas the tumour suppressor gene *Hao2* was upregulated (**Supplementary Fig. 4b**). Consistent with the PTPRK-ICD interactome results, cap-dependent translation initiation pathway was enriched in the Reactome GSEA pathway analysis (**Supplementary Fig. 4c**). Transforming growth factor beta receptor superfamily signalling and extracellular matrix organization pathways were suppressed in *Ptprk^ΔHep^* liver tumours in the GO GSEA analysis (**Fig. 3f**). Together with the suppressed activity of classical cancer pathways, including Ras signalling and MAPK signaling, observed in the KEGG GSEA analysis (**Fig. 3g**), these results suggest that hepatocyte PTPRK deletion suppressed, at least in part, obesity-associated HCC initiation.

Taken together, these results confirm and extend our previous findings, showing that hepatocyte-specific PTPRK deletion mitigates malignant cell transformation in the context of obesity-associated HCC. Our *in vivo* data provide evidence that hepatocyte PTPRK acts upstream of the PPARγ signalling network, promoting PPARγ-driven lipogenesis and steatotic adaptation during metabolic stress. The consistency of the *Ptprk^ΔHep^*phenotype with that of *Ptprk^-/-^* mice reinforces the conclusion that improved steatosis and reduced tumour initiation are primarily due to loss of PTPRK function in hepatocytes.

## Discussion

This study establishes that hepatocyte PTPRK is largely dispensable for metabolic homeostasis under chow feeding but becomes functionally important when nutrient excess engages steatotic adaptation. Using *Ptprk^ΔHep^* mice, we showed that hepatocyte PTPRK is sufficient to promote steatosis, insulin resistance, and increased tumour initiation under an obesogenic diet. The results observed in this study strengthen the notion that hepatocyte PTPRK acts upstream of obesity-induced PPARγ-linked lipogenesis and insulin resistance, shaping tumour susceptibility in a context of carcinogen exposure and hypercaloric diet feeding.

At the cellular level, estimation of cell size based on cortical actin staining showed that hepatocytes isolated from obese *Ptprk^ΔHep^*mice display reduced size, consistent with impaired steatosis-associated hypertrophy. This phenotype could reflect diminished PPARγ-driven lipogenesis,^11^ a pathway central to hepatocyte enlargement and lipid storage under nutrient excess. PTPRK is a membrane-bound phosphatase capable of homophilic dimerization and modulation of cell-cell adhesion, features that may influence actin cytoskeleton architecture and cellular expansion during metabolic stress.^21,22^ Structural studies of PTPRK and its close homologs support a capacity to integrate signalling and physical interactions at the plasma membrane, with potential consequences for lipid storage and compartmentalisation under obesogenic conditions.^22,23^

Our interactome findings reveal that PTPRK engages a hepatocyte-specific protein network distinct from that described in epithelial systems. ^24^ Previous work demonstrated that PTPRK displays marked substrate selectivity in epithelial cells, where it directly dephosphorylates junctional proteins such as Afadin, PARD3, p120-catenin, and δ-catenins, thereby preserving epithelial integrity and limiting invasive behaviour.^24^ In contrast, in primary mouse hepatocytes under conditions of induced tyrosine hyperphosphorylation, PTPRK associates with proteins enriched in metabolic, translational and trafficking functions. Thus, the downstream interactome and functional outputs of PTPRK seem to be cell-type dependent, supporting junctional regulation in epithelial cells^24^ and metabolic coordination in hepatocytes. However, transcriptome analysis of liver tumours showed reduced GO enrichment for cell junction assembly and regulation of cell-cell adhesion, while KEGG analysis showed enrichment of focal adhesion pathways in *Ptprk^ΔHep^* tumours. PTPRK has also been recently implicated in non-hepatic stress responses, promoting inflammation and pain-associated behaviours through a DUSP1-p38 MAPK axis in dorsal root ganglia.^25^ These studies reinforce the concept that PTPRK functions as a context-dependent signalling protein capable of coupling cellular stress to downstream adaptive programs.

PTPRK contains an inactive D2 pseudo-phosphatase domain, enabling recruitment of proteins independently of enzymatic action.^26^ Trapping mutants stabilise enzyme-protein complexes by arresting distinct steps of the phosphatase reaction cycle, allowing interactions that would normally be transient. We observed that many proteins associate with PTPRK independently of catalytic activity. This supports a scaffold-like role for PTPRK in organising metabolic protein assemblies. Among the proteins preferentially enriched in the PTPRK trapping mutants, Rab10 regulates vesicular trafficking, nutrient transporter recycling, and metabolic control.^27,28^ It has been shown to control endosomal recycling of glucose and lipoprotein receptors in hepatocytes and has been implicated in glucose uptake, lipophagy, and cholesterol handling, processes that directly influence hepatocarcinogenesis.^29^

Galactokinase 1 (GALK1), which catalyses the first committed step of galactose metabolism, was also enriched in the PTPRK trapping mutants. Perturbations in galactose metabolism are a cause of early-onset cataracts^30^, a phenotype previously reported in *Ptprk* knockout mice (The Jackson Laboratory). These observations suggested a broad role for PTPRK in carbohydrate metabolism, upstream of glycolysis and lipogenesis. In addition, the PTPRK-associated protein Slc2a9 (glucose transporter 9), a solute carrier involved in monosaccharide and organic anion transport,^31^ suggest that PTPRK-associated complexes converge on the regulation of carbon entry, intracellular trafficking, and hexoses metabolism. Defining whether these proteins are directly regulated by tyrosine phosphorylation, or instead reflect scaffolded assemblies stabilised by PTPRK under conditions of elevated phosphotyrosine signalling, will require targeted phosphosite mapping and functional validation.

DEN-treated obese *Ptprk^ΔHep^* mice developed significantly fewer tumours than controls, but with comparable liver and body weights, indicating that tumorigenic differences were not due to systemic weight changes. This supports a hepatocyte intrinsic requirement for PTPRK in liver tumour development. The hepatocyte-intrinsic role promoting HCC contrasts with the tumour-suppressive function previously described for PTPRK in epithelial tissues, where its primary activity restrains proliferation through junctional regulation and maintenance of tissue architecture.^16,21,32^ Our studies showed that PTPRK sustains a glycolytic and lipogenic state that buffers nutrient overload, while creating a metabolic environment permissive for carcinogen-induced transformation.

In conclusion, our findings identify hepatocyte PTPRK as a central mediator of steatotic adaptation, insulin resistance, and tumour initiation under metabolic stress. PTPRK is a potential target for strategies aimed at reversing MASLD and reducing liver cancer risk.^8^

## Materials and Methods

### Mouse and Metabolic Analysis

All animal experiments were conducted in compliance with Belgian regulations for animal welfare and approved by the Commission d’Éthique du Bien-Être Animal (protocol numbers 732 and 917). Mice were housed in the animal facility of Université libre de Bruxelles under controlled conditions (12-hour light/dark cycle, 22 °C) with ad libitum access to food and water.

*Ptprk* floxed (*Ptprk^Lox/Lox^*) mice were generated by homologous recombination in embryonic stem cells on a C57BL/6N background, with loxP sites inserted to flank exon 3 of the *Ptprk* gene. Targeting vector design, ES cell targeting, clone selection, and chimaera production were performed by Taconic Biosciences (TKC-210906-CAA-01-TAC). Targeted ES cell clone 1B2 was used to generate chimeric mice, which were bred to C57BL/6N mice to establish germline transmission. Heterozygous founders were intercrossed to obtain homozygous *Ptprk^Lox/Lox^*mice.

For hepatocyte-specific deletion, *Ptprk^Lox/Lox^* mice were crossed with Alb-Cre transgenic mice expressing Cre recombinase under the albumin promoter (Jackson Laboratory, RRID: IMSR_JAX:003574), generating *Ptprk^ΔHep^*mice and Cre-negative *Ptprk^Lox/Lox^* littermate controls. Male mice were used for the study and maintained under standard housing conditions and fed a standard chow diet (RM1 (P) 801151, Special Diets Services, UK) unless otherwise specified. At 8 weeks of age, mice were randomised to remain on chow or to receive a high-fat, high-fructose, high-cholesterol diet (HFHFHCD; D09100310i, Research Diets; 40% kcal from fat, 20% kcal from fructose, 2% cholesterol) for the indicated durations.

Body composition and liver fat/lean mass were assessed using EchoMRI^TM^ (Echo Medical Systems, Houston, TX, USA). Metabolic parameters, including energy expenditure, respiratory exchange ratio (RER), and physical activity, were measured using the TSE PhenoMaster system (TSE Systems, Germany). All measurements and endpoint analyses followed our described protocols.^11^

### Hyperinsulinemic-euglycemic Clamp

Hyperinsulinemic-euglycemic clamp experiments were performed in *Ptprk^ΔHep^*and control male mice.^33,34^ Briefly, 14-week-old male mice fed an HFHFHCD from 8 to 14 weeks of age were surgically implanted with a catheter in the right jugular vein. A single-channel vascular access port (VABM1B/25, Instech Laboratories) was exteriorised at the back of the neck using a magnetic button system. Mice were allowed to recover for 10 days post-surgery before undergoing clamp procedures. During the clamp, insulin (Actrapid®, 100 U/mL; Novo Nordisk) was infused continuously at a rate of 2 mIU/kg/min for HFHFHCD-fed mice. Blood glucose levels were monitored at regular intervals using a StatStrip® Xpress2™ glucometer (Nova Biomedical, #56506) to maintain euglycemia.

### DEN-Induced Hepatocellular Carcinoma (HCC)

Hepatocellular carcinoma was induced by a single intraperitoneal injection of DEN at a dose of 25 mg/kg body weight, diluted in PBS, administered to 2-week-old male mice. Starting at 6 weeks of age, mice were fed an HFHFHCD to promote obesity-associated liver tumorigenesis. Mice were sacrificed at the experimental endpoint by cervical dislocation. Livers were harvested and subjected to comprehensive macroscopic and histological evaluation to assess tumour number and size.

### CT Scan and Reconstruction

High-resolution micro-computed tomography (µCT HR) imaging was performed *in vivo* to assess liver tumour burden. Mice received an intravenous injection (tail vein) of Exitron™ nano 12000 (Miltenyi Biotec) prior to scanning to enhance hepatic tissue contrast. Image acquisition was carried out under anaesthesia using a µCT scanner (Skyscan 1276, Brucker), focusing on the abdominal region. Acquisition of around 7min duration was done with the X-rays source put at 90 kV, 200 µA with a 0.5 mm Al filter in a step and shot scanning mode of 360 degrees, a rotation step of 0.8-degree, exposure time of 99 ms per step and an averaging of 2 frames, binning 4 x 4 to provide images of 1008 x 672 pixels of 40.9 µm. Flat Field Correction is applied. Images were reconstructed with NRecon software v2.2.0.6 (Brucker) with a CS to Image Conversion of 0.000832 set as minimum and 0.033000 set as maximum, a smoothing of 1, a Beam Hardening Correction of 0% and a ring artefact correction is adjusted individually for each sample. Tumour detection was done by a visual, qualitative analysis of each frame following image reconstructions.

### Histological Analysis

Liver and tumour tissues from euthanised control and *Ptprk^ΔHep^*mice were dissected, fixed in 4% buffered formaldehyde (pH 7.4), and embedded in paraffin for histological evaluation. Sections of 7 μm thickness were prepared using a Leica rotary microtome and stained with H&E to assess liver architecture and tumour morphology. Whole-slide images were acquired at 40X magnification using the NanoZoomer Digital Pathology system (Hamamatsu Photonics K.K., version SQ 1.0.9).

Mouse livers were collected and fixed overnight in 4% paraformaldehyde (PFA) at 4°C, followed by cryoprotection in 30% sucrose at 4°C until the tissues sank. Samples were then embedded in Tissue Freezing Medium (Leica, 14020108926), frozen, and stored at -80°C. Liver blocks were sectioned at 50 µm using a cryostat (Leica CM3050 S) and mounted on adhesion slides (Epredia, J1830AMNZ). Sections were air-dried at room temperature for 1-2 hours, then washed with PBS to remove residual embedding medium. Hydrophobic barriers were drawn around the tissue using a liquid-repellent slide marker pen. For staining, Phalloidin-iFluor™ 633 Conjugate (AAT Bioquest, 23125) was reconstituted in 30 µL DMSO and diluted 1:300 in PBS to prepare the working solution. Sections were incubated with the phalloidin solution overnight at 4°C, followed by three washes with 1× PBS. Nuclear staining was performed using 1 µg/mL DAPI (Carl Roth, 28718-90-3) for 5 minutes at room temperature, followed by three PBS washes. Slides were mounted with Fluoromount-G (ThermoFisher, 00-4958-02) and imaged using a confocal microscope (LSM 780, AxioObserver) equipped with a 20X objective (Plan-Apochromat 20×/0.8 M27). Laser settings were as follows: Phalloidin-iFluor™ 633: excitation at 633 nm (12%), detection at 643-755 nm; DAPI: excitation at 405 nm (3%), detection at 410-471 nm. Image analysis and hepatocyte surface quantification were performed using the “Cell” module in Imaris software.

### Western Blotting

Total protein lysates were extracted from liver tissues using RIPA buffer (Cell Signalling Technology) supplemented with Halt™ protease and phosphatase inhibitor cocktail (Thermo Fisher Scientific). Protein concentrations were quantified using the BCA Protein Assay Kit (Thermo Fisher Scientific).

For immunoblotting, 20-30 μg of protein was resolved on polyacrylamide gels and transferred to 0.22 μm nitrocellulose membranes (Bio-Rad Laboratories). Membranes were blocked with 5% non-fat milk in TBST and incubated overnight with primary antibodies (listed in **Supplementary Table S1**) diluted in 5% BSA/TBST. The following secondary antibodies were used: goat anti-mouse IgG (Dako Agilent), goat anti-rabbit IgG (Dako Agilent), and Peroxidase AffiniPure donkey anti-human IgG (Jackson ImmunoResearch). Immunoreactive bands were visualised using the Amersham ImageQuant 800 imaging system (Cytiva Life Sciences).

### RNA Extraction, RT-PCR, and Bulk RNA Sequencing

Total RNA was isolated from liver tissues or tumour tissues using the RNeasy Mini Kit (QIAGEN) following the manufacturer’s protocol. Liver tissue cDNA synthesis was performed using a reverse transcriptase kit (Eurogentec). Quantitative real-time PCR (qRT-PCR) was conducted using SYBR Green reagents (Bio-Rad Laboratories) on a Bio-Rad CFX system. The procedure followed previously described methods ^11^. Primer sequences are provided in **Supplementary Table S2**. Total RNA quality analysis, library preparation (poly A enrichment), and sequencing were performed by Novogene (UK). Differential gene expression analysis was performed using DESeq2 and gene set enrichment analysis (GSEA) was performed using fGSEA, which were described previously.^34^

### Mass Spectrometry-Based Interactome Analysis

Interactome profiling was performed in primary mouse hepatocytes for four experimental conditions: wild-type (WT, control for basal expression), cysteine-serine mutation (CS), aspartate-alanine mutation (DA), and beads-only (negative control). Affinity purification was conducted on cell lysates to capture interactors using recombinant proteins or beads alone. Protein eluates were analysed by liquid chromatography-tandem mass spectrometry (LC-MS/MS). Protein eluates were run 5-10mm into SDS-PAGE gels and stained with Coomassie. The protein-containing gel regions were reduced and carbamidomethylated, then digested with trypsin (Promega) at 30°C overnight. Tryptic peptides were analysed by nanoLC-MS/MS on Orbitrap Eclipse Tribrid mass spectrometer coupled to an UltiMate 3000 RSLCNano (ThermoFisher). The peptides were separated on a PepMap C18 column (75 µm id x 50 cm, 3 µm beads, ThermoFisher) with a linear 60 min gradient from 2% to 40% acetonitrile/0.1%FA, at a flow rate of 300 nl/min. The MS1 survey scan range was m/z 300 – 1500 at a resolution of 120,000, AGC at 4 x 10^5^ and maximum IT at 50 msec. Multiply charged precursors (from 2+ to 7+) were selected for HCD fragmentation with an isolation width at 0.7 Th, with a 2 sec cycle time. The intensity threshold was set at 5 x 10^4^, the collision energy at 30%, AGC at 1×10^4^, max IT at 200 ms and the resolution at 15,000. The dynamic exclusion time was set at 60 sec with ± 10 ppm mass tolerance.

The raw files were processed in Proteome Discoverer 2.5 (ThermoFisher) using the Sequest HT search engine to search against the Uniprot human canonical proteome database and a database of common contaminants. Trypsin was set at full specificity with a maximum of two miss-cleavage sites. The mass tolerances were set at 10 ppm for the precursors, and 0.02 Da for the fragment ions. Oxidation (M) and acetylation/Met-loss (protein N-terminus) were set as dynamic modifications, and carbamidomethyl (C) as static modification. Peptides were validated by Percolator with q-value of 0.01 for the decoy database search, and only high confident PSMs (Peptide Spectrum Matches) were considered. Protein FDR was set at 0.05. Only master proteins were reported.

^24^Protein intensity matrices generated in PD were analysed in R (v4.1.2). Proteins lacking unique identifiers, with incomplete quantification, or uncertain gene mapping were excluded. Protein intensities were normalised and log₂-transformed using the Voom method from the limma package (v3.50.1). Principal component analysis (PCA) was used to assess batch effects and data structure. Differential abundance analysis was performed using limma with empirical Bayes moderation. Pairwise contrasts were computed for all conditions, applying a significant threshold of *p* < 0.05 and |log₂(fold change)| > 0.5. Volcano plots were generated using MagmaFlow (v10.0.3, https://zenodo.org/records/17107683).

Over-representation analysis (ORA) was performed using curated gene sets from the Kyoto Encyclopedia of Genes and Genomes (KEGG), Gene Ontology (GO), and Reactome. Analyses were conducted in R using the clusterProfiler and enrichR packages. For CS vs. WT and DA vs. WT comparisons, both upregulated and downregulated proteins were used as input gene sets. For comparisons against beads-only, only upregulated proteins were included. For glycolysis pathway analysis, all significant proteins identified as PTPRK interactors in any comparison were included, regardless of regulation direction. This approach enabled a comprehensive assessment of PTPRK-associated glycolytic regulation. Enrichment results were visualised with dot plots using ggplot2. Heatmaps of variably enriched proteins were generated with pheatmap. Venn diagrams were created using matplotlib-venn in Python.

## Statistical Analysis

Sample size (n) represents individual mice or independent experimental replicates. Data are presented as mean ± standard error of the mean (SEM). Statistical analyses were performed using GraphPad Prism v10 (GraphPad Software, La Jolla, CA, USA). Group comparisons were conducted using Student’s *t*-test, one-way ANOVA, or two-way ANOVA, as appropriate. Sample sizes were determined based on preliminary data and variability from previous studies.^11^ Statistical significance was defined as follows: * *p* < 0.05, ** *p* < 0.01, *** *p* < 0.001.

## Acknowledgement

We thank Anne Van Praet, Mariana Nunes, Francisco Costa, and Cláudia S. Pinto (Université libre de Bruxelles) for experimental and technical support. We thank Latifa Bakiri (Medical University of Vienna) for critical reading of the manuscript. We have used BioRender.com for the design of cartoons in the different panels (under license). The European Research Council (ERC) Consolidator grant METAPTPs grant Agreement No. GA817940 (ENG); FNRS-PDR grant 40020272 (ENG); FNRS-TELEVIE grants 40007402, 40018756, 40025595 (ENG); Fonds Paul GENICOT (ENG); The ULB Foundation (ENG); PhD scholarship support from China Scholarship Council (AL); SPS is supported by Ramalingaswami Re-entry Fellowship from the Department of Biotechnology (DBT), India; ENG is a Research Associate of the FNRS, Belgium.

## Data Availability

Enrichment results are available in the Supplementary Excel files. Custom R scripts (https://github.com/carlosbuss1/PTPRK_Interactome_Project). Raw mass spectrometry data and processed interactome datasets are available upon reasonable request.

