## Supplementary Material for "Loss of PTPRK in hepatocytes reduces steatosis and carcinogen-induced tumour development in obesity"

| Primary antibody | Designation | Source or reference | Cat# | Additional information |
|---|---|---|---|---|
| PTPRK | Human monoclonal anti-PTPRK | Feamley et al. 2019, Kindly provided by Hayley Sharpe | H4 | WB 1:2000 |
| PPARγ | Rabbit monoclonal anti-PPARγ | Cell Signalling Technology | 2443 | WB 1:500 |
| β-actin | Anti-β-Actin antibody, Mouse monoclonal | Sigma-Aldrich | A1978 | WB 1:2000 |

**Table S1.** List of primary antibodies used for Western blotting.

| Gene | qPCR F | qPCR R | Standard F | Standard R |
|---|---|---|---|---|
| *Gapdh* | AGTTCAACGGCACAGTCAAG | TACTCAGCACCAGCATCACC | ATGACTCTACCCACGGCAAG | TGTGAGGGAGATGCTCAGTG |
| *Pparγ1* | CCAAGAATACCAAAGTGCGATCA | AAAACCCTTGCATCCTTCACAA | GCTCCAAGAATACCAAAGTGCGA | AACCTGATGGCATTGTGAGACA |
| *Pparγ2* | TGCCTATGAGCACTTCACAAG | TCTACTTTGATCGCACTTTGGTA | AGCATGGTGCCTTCGCTGAT | GCCCAAACCTGATGGCATTGTG |

**Table S2.** List of primers used for RT-PCR.

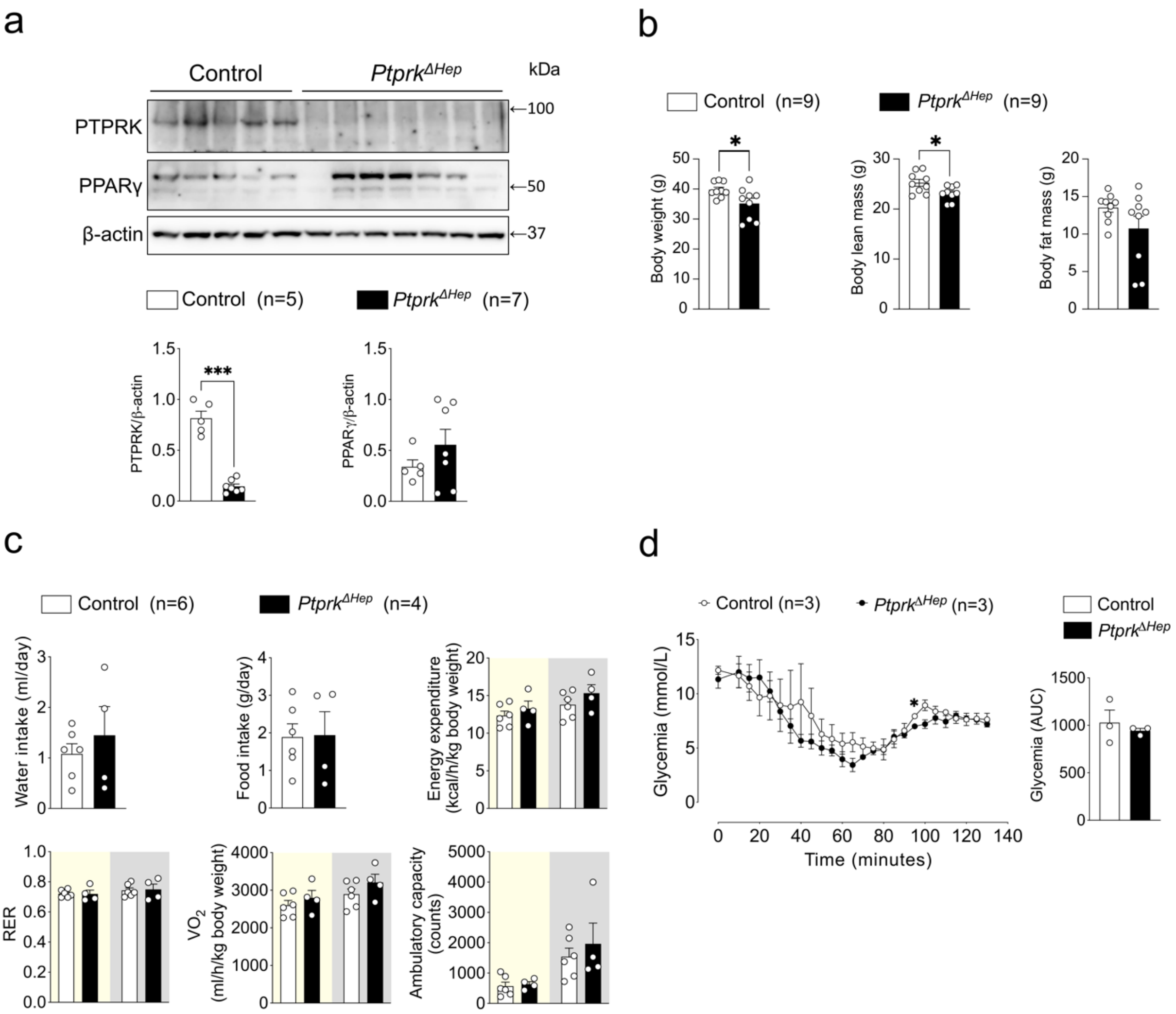


**Supplementary Fig. 1. Immunoblot, body parameters, metabolic cage, and clamped glycemia analysis of liver-specific PTPRK-deficient and control.** (a) Immunoblot analysis of liver samples from *Ptprk*$^{\Delta Hep}$ and control mice fed a chow diet for 12 weeks (n=5-7). (b) Body weight and EchoMRI measurements in *Ptprk*$^{\Delta Hep}$ and control mice after 6 weeks of HFHFHCD feeding (n=9). (c) Daily water and food intake, energy expenditure, respiratory exchange ratio (RER=$VCO_2/VO_2$), oxygen consumption ($VO_2$), and ambulatory activity (n=4-6). (d) Hyperinsulinemic-euglycemic clamp analysis showing glycemia in *Ptprk*$^{\Delta Hep}$ (n=3) and control (n=3) mice fed with HFHFHCD for 6 weeks (14-week-old). Data are presented as mean ± SEM (a-d). Statistical analyses were performed using a two-tailed unpaired Student's *t*-test (a-d). *$p < 0.05$, and ***$p < 0.001$.

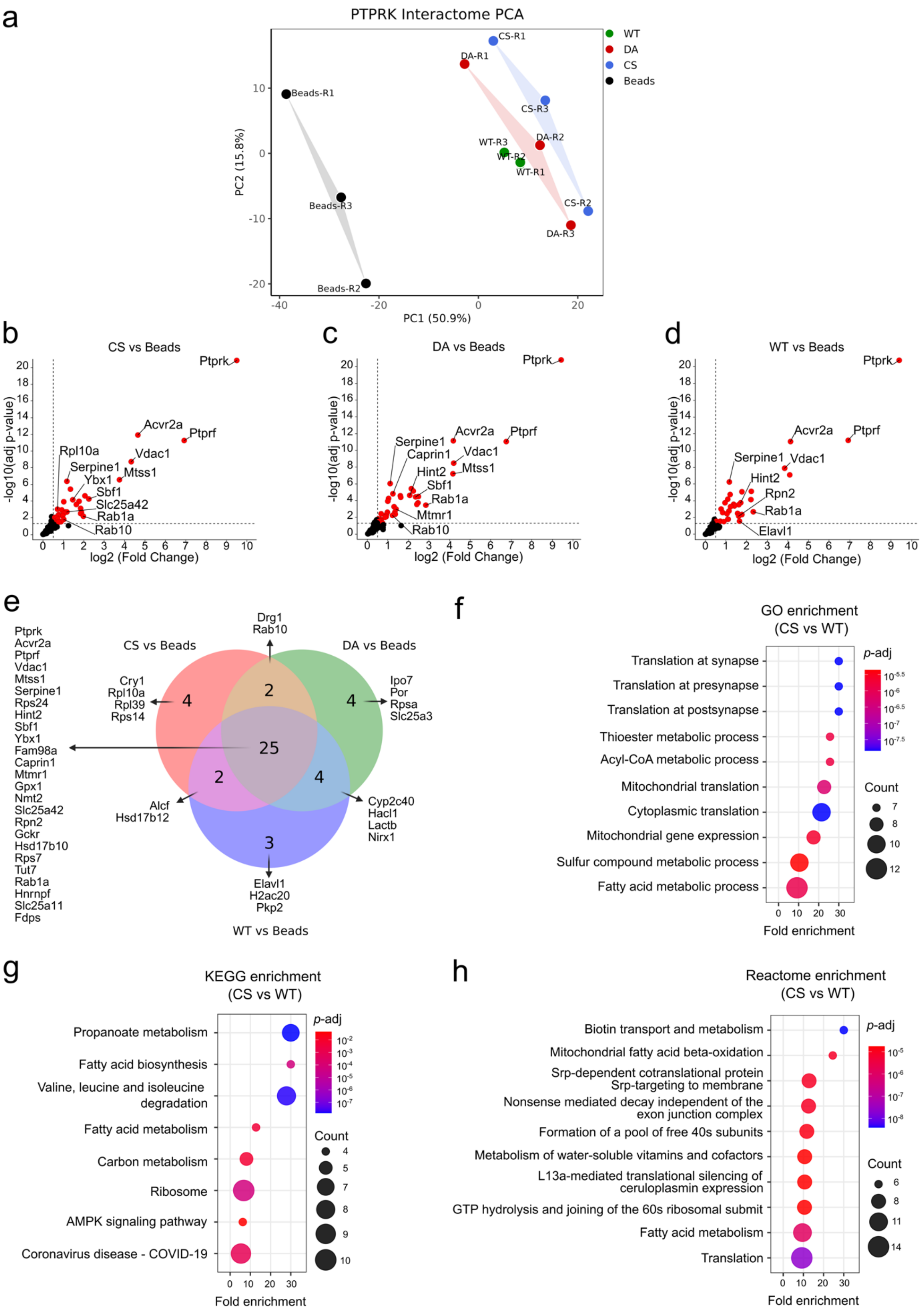


**Supplementary Fig. 2. PCA, volcano, Venn, and ORA pathway enrichment plots of PTPRK interactome.** (a) PCA plot of interactome in beads, WT, DA, CS groups (n=3). (b-d) Volcano plots of enriched proteins in CS vs Beads (b), DA vs Beads (c), and WT vs Beads (d). (e) Venn diagram of significantly enriched p-tyrosine pulldown proteins in WT, DA, and CS compared to beads. (f-h) ORA pathway enrichment analyses of interactome in CS vs WT, including GO (f), KEGG (g), and Reactome (h). In (f-h), FDR < 0.05.

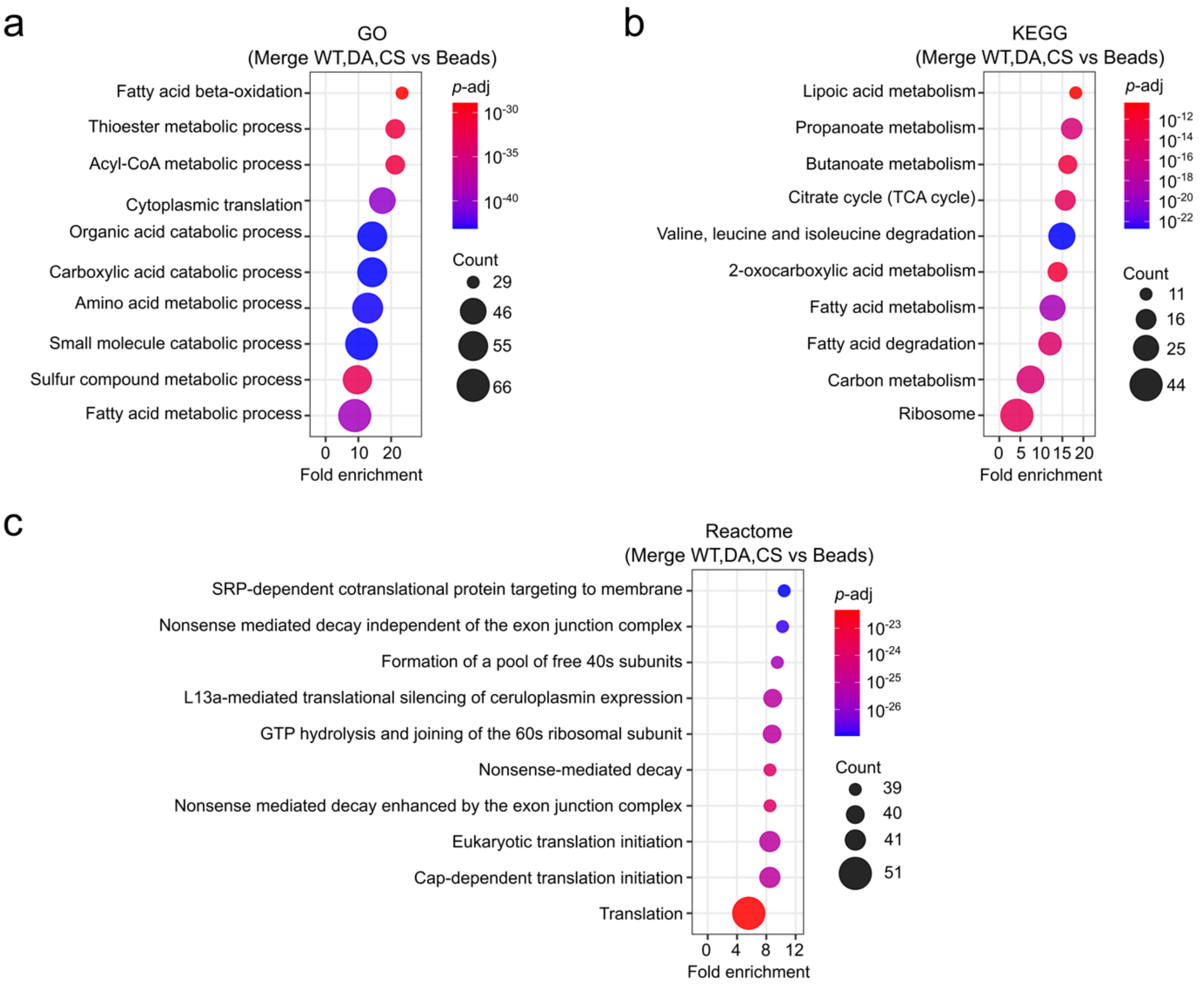


**Supplementary Fig. 3. ORA pathway enrichment analysis of interactome indicating the top-enriched pathways in GO, KEGG, and Reactome.** (a-c) ORA using all significantly enriched proteins binding to WT, DA, and CS, including GO (a), KEGG (b), and Reactome (c). The proteins that only bound to Beads were removed before the pathway analysis. In (a-c), FDR < 0.05.

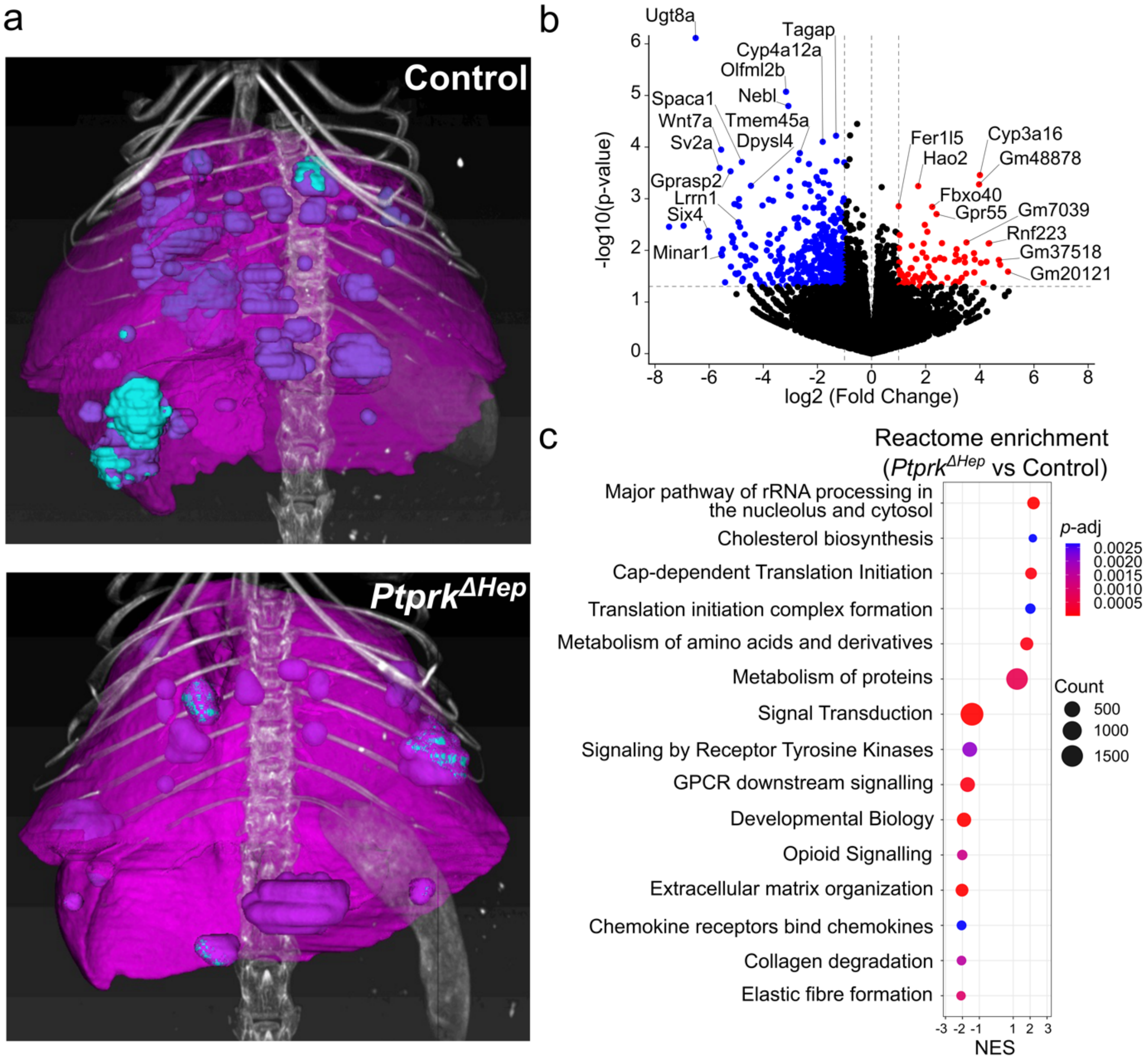


**Supplementary Fig. 4. Tumour 3D reconstructed images, differential gene expression and signalling pathways altered by PTPRK.** (a) Representative 3D reconstruction images of tumour development (blue-violet nodules) *Ptprk*$^{\Delta Hep}$ and control mice. (b) Volcano plot of differentially expressed genes in *Ptprk*$^{\Delta Hep}$ vs control tumours (n=6). (c) Reactome GSEA pathway enrichment analyses of transcriptomics in *Ptprk*$^{\Delta Hep}$ vs control. In (d), FDR < 0.05.
